# A new method and application of PacBio sequencing for low copy and difficulty preparation plasmids

**DOI:** 10.1101/2024.05.07.593053

**Authors:** Xiaoshu Ma, Yusha Wang, Ruikai Jia, Hua Ye

## Abstract

The Single Molecule Real Time (SMRT) system developed by Pacific Biosciences applies the principle of synthesis while sequencing and uses the SMRT chip as the sequencing carrier. The high starting amount and good integrity of DNA required by PacBio library construction has always been a headache. Generally, the total loading volume of PacBio sequencing is 10μg, and the concentration is not less than 200ng/μL, which is a huge challenge for low-copy plasmids. Tight plasmid means that when the bacterial chromosome replicates once, the plasmid replicates once, and each bacterium only contains 1 to 2 plasmids. The replicon of the plasmid determines the copy number of the plasmid. Low copy plasmids should produce 0.2-1 μg DNA per ml of LB culture. Low-copy plasmids play a key role in gene synthesis. When vectors are used for expression or other purposes, low-copy plasmids are used to reduce resource consumption caused by plasmid expansion. Low-copy plasmids are also used when plasmids have lethal gene clones. In order to solve the problem of low copy plasmid library database construction in PacBio, we used Phi29 polymerase to perform multiple substitution amplification of low copy plasmid, so as to obtain a large number of high molecular weight DNA to meet the computer requirements of PacBio. In addition, this study also established a PacBio sequencing method for bacterial fluids without the need for plasmid extraction steps, thereby reducing time and money costs.

## Introduction

Sequel, a third-generation sequencing system based on PacBio single molecule real-time sequencing technology (SMRT), is a relatively mature third-generation sequencing technology [1]. It has the characteristics of long read length, high consistent accuracy, single molecule sequencing, etc. At the same time, it avoids sequencing bias introduced by PCR and can obtain base modification information at the same time [2].

The Single Molecule Real Time (SMRT) system developed by Pacific Biosciences applies the principle of synthesis while sequencing and uses the SMRT chip as the sequencing carrier [3], which can provide longer read length and faster running speed. There are many ZMW (Zero-Mode Waveguides) holes in SMRT [4]. Each SMRT Cell contains a large number of ZMW circular nanoholes with a diameter of 50∼100nm, which uses a physical effect zero-mode waveguide with an outer diameter smaller than the wavelength of excited light [5]. When DNA molecules enter the hole, the excited light emitted from the bottom of the hole cannot penetrate the hole into the solution region above. It is confined only to an area at the bottom that is large enough to cover the part of the DNA being detected, thereby collecting signals from that area and minimizing background noise [6].

SMRTbell database library is a single strand hairpin adaptor [7], which can connect long DNA molecules with sequencing adaptor to form a stem ring structure, and the stem ring structure can be combined with sequencing primers and DNA polymerase molecules complementary to the adaptor to form a sequencing complex, which can be sequenced on PacBio RS.On-machine sequencing is to put the constructed library complex into the PacBio RS sequencer and load it into the nanopore of the sequencing chip SMRT Cell. DNA polymerase molecules are fixed at the bottom of the nanopore through covalent binding, usually one DNA molecule is fixed in each nanopore. dNTP, the substrate required for DNA polymerization, and buffer are added into the hole of the SMRT Cell chip. The four dNTPs have fluorescent labeling groups of four colors. According to the nucleotide sequence of the template chain, the corresponding dNTP enters the DNA template chain and reacts with the primer and polymerase complex [10]. At the same time, dNTP fluorescence signal was detected by zero-mode waveguide, fluorescence signal image was obtained, and DNA base sequence was obtained through calculation and analysis [11].

Plasmids are a class of nucleic acid molecules inherent in biological cells that can replicate autonomously and be stably inherited independently of host chromosomes [12].

Plasmids are commonly found in prokaryotic bacteria and fungi, and most of them are DNA-type, while a few are RNA type [13]. Most natural DNA plasmids have a covalent, closed, circular molecular structure, and their molecular weight ranges from 1 to 300 kb.Bacterial plasmids are the most commonly used carriers in genetic engineering. Plasmids can autonomously replicate by utilizing the host cell’s DNA replication system [14]. The replicator structure on their DNA determines the correspondence between the plasmid and the host, and determines the number of molecules (copy number) in each cell.Plasmids can be divided into two major replication types [15] : tight replication-controlled plasmids and relaxed replication-controlled plasmids.

Relaxed plasmid: When bacterial chromosome replication stops, it can continue to replicate, and each bacterium generally contains about 20 copies of the plasmid [16]. The replication of these plasmids is under the relaxed control of the host, and each bacterium contains 10-200 copies, which can be increased to several thousand copies if the host protein synthesis is inhibited by certain drug treatment [17]. Tight plasmid: When the bacterial chromosome replicates once, the plasmid replicates once, and each bacterium only contains 1 to 2 plasmids [18]. The replicon of the plasmid determines the copy number of the plasmid. The copy number of plasmids is related to the relative intensity of positive and negative regulation, and can also be changed by mutating replicons. There are two main uses of low copy number plasmids: (1) to enlarge the fragment or to have lethal gene cloning in the amplification of high copy number plasmids [19]. 2. The expansion of plasmids will occupy a lot of resources, and low-copy plasmids will also be used when vectors are used for expression or other purposes [20]. Low copy plasmids produce only 0.2-1 μg DNA per ml of LB culture [21]. However, the DNA required for the construction of the Pacbio sequencing library has a high starting amount and good integrity, and the total amount of samples for PacBio sequencing is 10μg, the precious samples are not less than 5μg, and the concentration is not less than 200ng/μL. The minimum amount and concentration are subject to the Qubit detection results. Compared with nanodrop, the detection results are more stringent. All of these make it difficult for low-copy plasmids to meet the sequencing needs, and the challenge is huge.

Multiple displacement amplification (MDA) is isothermal chain displacement amplification [22]. Under constant temperature, a random primer composed of 6 random bases is randomly annealed with the template, followed by a chain displacement reaction under the action of phi29 DNA polymerase. After replacement, the single strand can randomly bind, anneal and extend with the primers, and finally form branch amplification. Compared with the WGA method of PCR [23], the bias is 3 times smaller [24], and MDA can produce about 20 to 30 μg of product from 1 to 10 DNA copy amplification [25].

Phi29 polymerase has strong chain displacement activity, which can realize the unchain and replication of the complex structure of DNA, and has strong chain affinity. A single polymerization reaction can achieve continuous polymerization extension of up to 100 kb [26]. In addition, Phi29 polymerase also has strong 3 ’-5’ outcut collation activity [27]. Its fidelity is 1000 times that of Taq enzymes, which is higher than the fidelity of most current high-fidelity enzymes, which guarantees high fidelity during DNA amplification [28]. In this paper, we will use Phi29 polymerase to perform multiple displacement amplification of low-copy plasmids, so as to obtain a large amount of high-quality DNA to meet the requirements of on-machine sequencing by PacBio.

## Result

### 1. After MDA, PacBio sequencing coverage increased significantly

We first selected two normal plasmids for pacbio sequencing. The original concentrations of the two plasmids were 209ng/μL and 95 ng/μL, but the concentrations of the plasmids were only 3.5ng /μL and 4.8ng /μL after library construction. After the construction of the database, the concentration is low, which has a great influence on the output of the subsequent data volume.

In view of this, we first carried out MDA reaction on these two plasmids, and then measured the concentration with Qubit, and the amount of plasmids increased by 10 times (Table 1). We can see from Table 2 that after MDA, the concentration of plasmids increased greatly, and the amount of data also increased greatly after pacbio was installed. The number of “X1” plasmid data increased from 43reads to 533reads, of which 487 matched the reference sequence, accounting for 91.37% of the total reads. The number of “X2” plasmid data increased from the initial 24reads to 605reads, of which 560 matched the reference sequence, accounting for 92.56% of the total reads. The increase in the amount of data can make our data more convincing. It can be seen from the sequencing results that the sequencing sequences of the two plasmids of “X1-MDA” and “X2-MDA” are consistent with the reference sequences after analysis (Figure 1).

**Figure 1.**
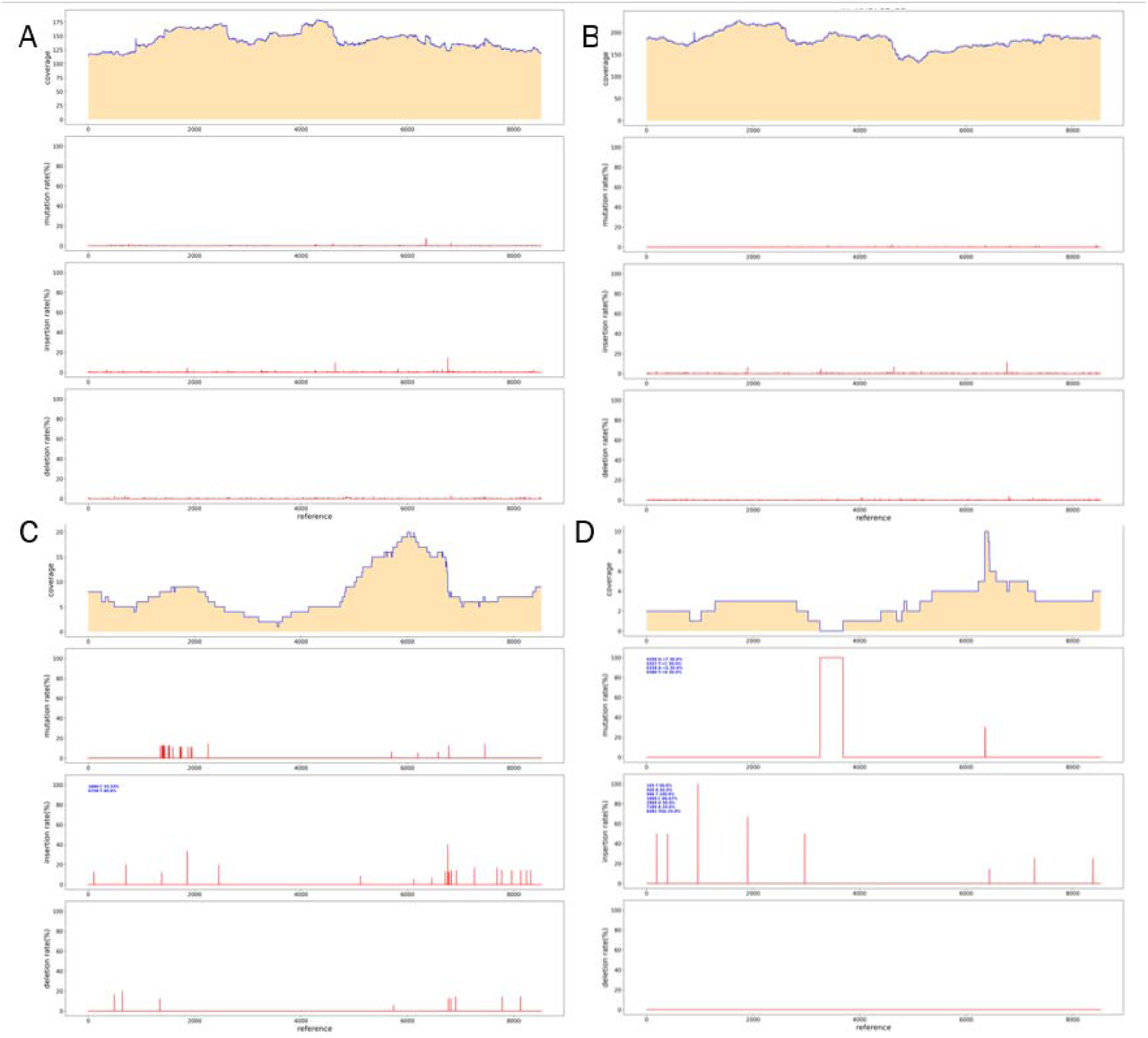
After rolling out replication, PacBio sequencing results. A. Results of PacBio sequencing of X1-MDA plasmid. B. results of PacBio sequencing of X2-MDA plasmid. C. Results of PacBio sequencing of X1 plasmid. D. results of PacBio sequencing of X2 plasmid.

**Table 1.** Library construction and plasmid concentration after MDA reaction.

| Sample Name | Original Concentration | Library Concentration ( 10 μL ) | Post- MDA Concentration ( 70μL ) | Post- MDA Library Concentration ( 10μL ) |
| --- | --- | --- | --- | --- |
| X1 | 95ng/μL ( 6μL ) | 3.5ng/μL | 200ng/μL | 6.8ng/μL |
| X2 | 209ng/μL ( 6μL ) | 4.8ng/μL | 210ng/μL | 10.2ng/μL |

**Table 2.** The amount of PacBio data used for database construction after MDA reaction.

| Sample Name | Total Reads | Mapped Reads | Percent (%) | Unmapped Reads | Percent (%) |
| --- | --- | --- | --- | --- | --- |
| X1 | 43 | 36 | 83.72 | 7 | 16.28 |
| X1-MDA | 533 | 487 | 91.37 | 46 | 8.63 |
| X2 | 24 | 15 | 62.5 | 9 | 37.5 |
| X2-MDA | 605 | 560 | 92.56 | 45 | 7.44 |

### 2. Comparison of the data volume output after plasmid MDA after database construction and before database construction

According to the experimental results in Figure 1, we speculated that the concentration level after library construction may be the key to directly affect the output of data volume after pacbio sequencing. As to whether the data volume can be increased directly by increasing the concentration of DNA library after library construction, without being limited to the step of low or high copy plasmid for MDA reaction, that is, whether the number of copies of the plasmid is or not, We tested our conjecture by direct MDA reaction only on the low concentration DNA library after the construction of the library.Therefore, we selected two plasmid libraries with low concentration after the construction of the library for MDA reaction, and the concentration of the library increased greatly after the reaction, from the original concentration of 1.4ng/μL to 200 ng/μL (Table 3). The products after MDA reaction were sequenced by pacbio, but the results were beyond our expectation.The library products after MDA reaction do not have any data volume, while the products without MDA reaction have a small amount of data output (Table 4). For this reason, we speculate that the template of the Pacbio library is a dumbbell shape, and the entire molecule is actually a ring. In the process of sequencing, it can be sequenced repeatedly, which is not only conducive to the advantage of PacBio’s long read length, but also conducive to multiple passes to correct random error rates. MDA uses random primers (typically 6bp) that can be annealed with template DNA at multiple sites, and then polymerase is used to start replication at multiple sites of DNA simultaneously, thus synthesizing complementary DNA along the DNA template. The MDA reaction damaged the circular library of pacbio, resulting in the suspension of circular sequencing and no data production. Therefore, if the problem of low DNA library after library construction is encountered, no matter whether the plasmid has high copy or not, MDA reaction should be performed on the plasmid before library construction to improve its data output.

**Table 3.** Library construction and plasmid concentration after MDA reaction.

| Sample Name | Original Concentration | Library Concentration ( 10μL ) | Post-MDA Library Concentration ( 10μL ) |
| --- | --- | --- | --- |
| X3 | 190ng/μL ( 3μL ) | 1.4ng/μL | 200ng/μL |
| X4 | 108ng/μL ( 3μL ) | 0.6ng/μL | 37.5ng/μL |

**Table 4.** The amount of PacBio data used for database construction after MDA reaction.

| Sample Name | Total Reads | Mapped Reads | Percent (%) | Unmapped Reads | Percent (%) |
| --- | --- | --- | --- | --- | --- |
| X3 | 9 | 4 | 44.44 | 5 | 55.56 |
| X3-TP | 0 | 0 | 100 | 0 | 0 |
| X4 | 9 | 8 | 88.89 | 1 | 11.11 |
| X4-TP | 0 | 0 | 100 | 0 | 0 |

### 3. Sequencing results of PacBio after MDA for low-copy plasmids with special structures

In order to further verify whether the MAD reaction is suitable for low-copy plasmids containing special structures, we loaded a fragment containing a large number of reverse repeats into the vector (Figure 2) and conducted sanger sequencing, which was correct. We selected two clones, each of which took 2μL plasmid, and carried out MAD reaction with phi29 enzyme. After the reaction, the concentrations were 216ng/μL and 213 ng/μL (Table 5). After the establishment of the library, the concentrations were 3.2 ng/μL and 3.5 ng/μL, and pacbio sequencing was performed. The total number of “X5” plasmid data was 3860reads, among which 2581 matched the reference sequence, accounting for 66.87% of the total reads. The total number of “X6” plasmid data was 2192reads, among which 1127 matched the reference sequence, accounting for 51.41% of the total reads (Table 6). From the analysis of reads numbers, these data volumes were sufficient to meet our requirements for plasmid analysis. After data analysis, it was found that the two samples “X5” and “X6” had deletion at 6984 bits, which was verified by sanger sequencing. It can be found that the sequencing result of sanger also had deletion at 6984 bits, which matched the sequencing result of pacbio (Figure 3). Therefore, we found that the complex structure of plasmids had no effect on MDA response.

**Figure 2.**
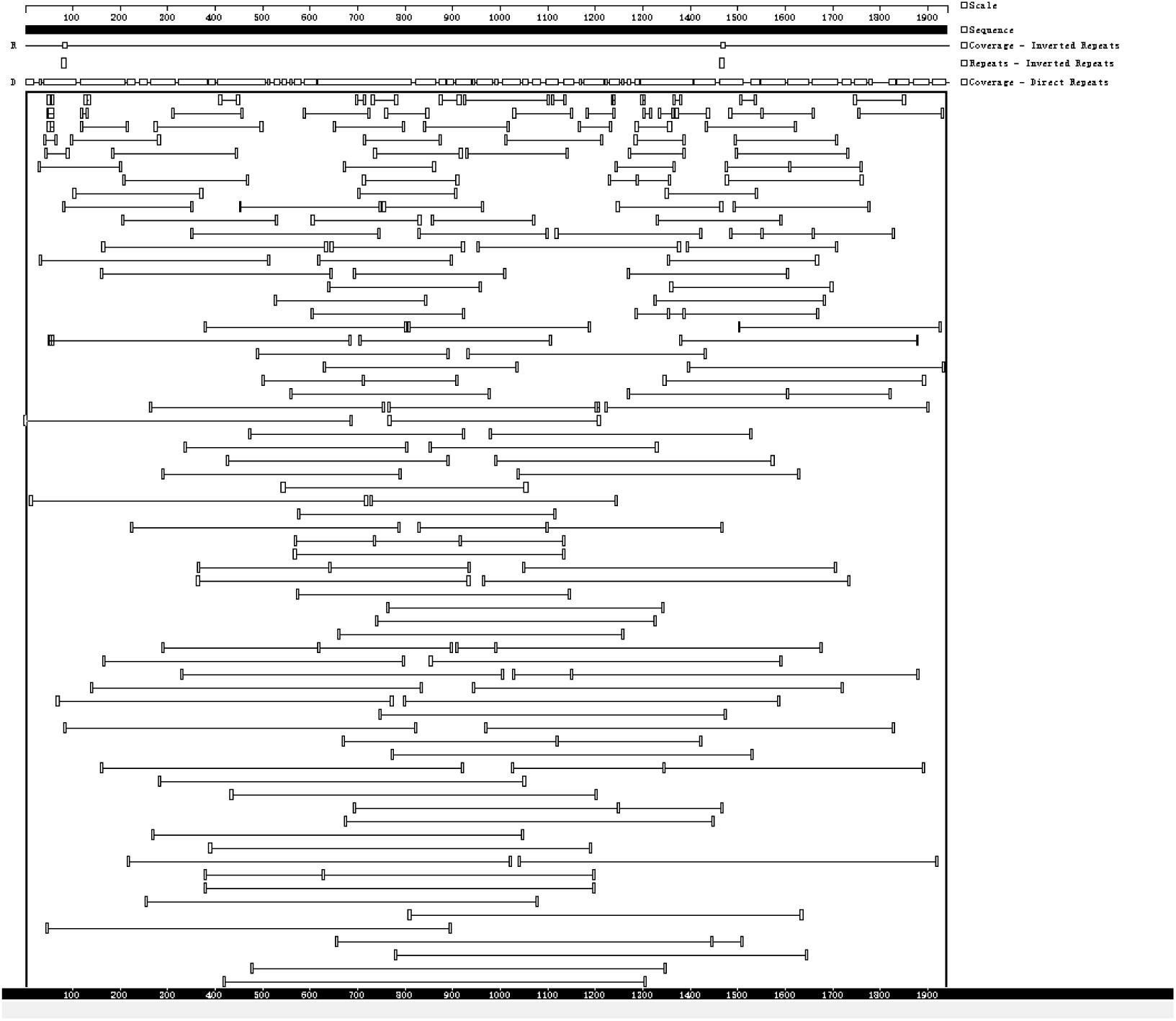
Sequence structure analysis of plasmids. The sequence structure was analyzed by Gene Quest software.

**Figure 3.**
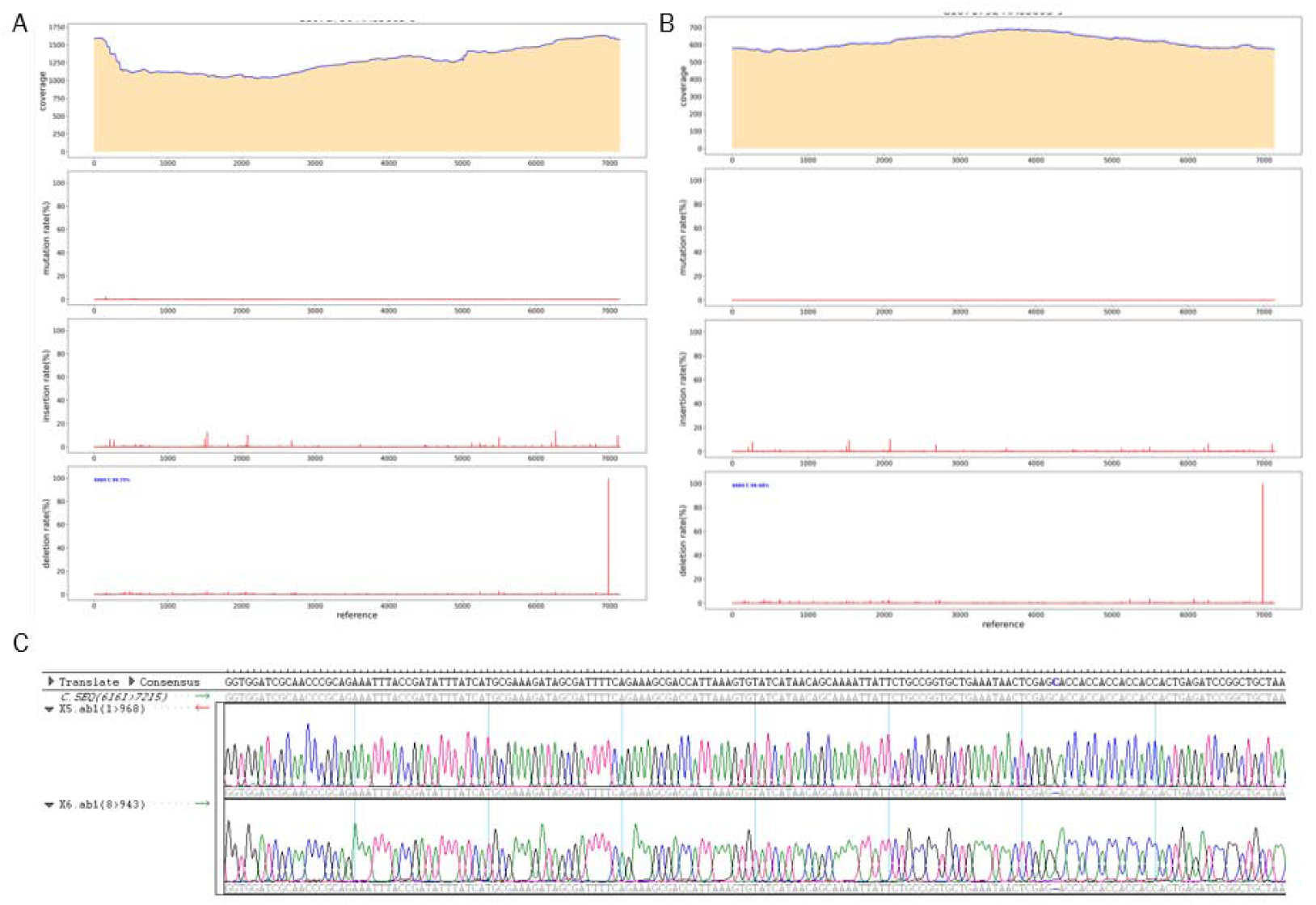
After rolling out replication, PacBio sequencing results. A. Results of PacBio sequencing of X5 plasmid. B. results of PacBio sequencing of X6 plasmid. C. Sanger sequencing results of X5 and X6 samples.

**Table 5.** Library construction and plasmid concentration after MDA reaction.

| Sample Name | Original Concentration | Post-MDA Concentration ( 10 μ L ) | Post- MDA Library Concentration ( 10μL ) |
| --- | --- | --- | --- |
| X5 | 55ng/μL ( 2μL ) | 216ng/μL | 3.2ng/μL |
| X6 | 48ng/μL ( 2μL ) | 213ng/μL | 3.5ng/μL |

**Table 6.** The amount of Pacbio data used for database construction after MDA reaction.

| Sample Name | Total Reads | Mapped Reads | Percent (%) | Unmapped Reads | Percent(%) |
| --- | --- | --- | --- | --- | --- |
| X5 | 3860 | 2581 | 66.87 | 1279 | 33.13 |
| X6 | 2192 | 1127 | 51.41 | 1065 | 48.59 |

### 4. Complex structure plasmid, sequencing results of PacBio after MDA

In order to further verify the effect of complex structure plasmids on MDA response. We constructed two plasmids with direct and reverse repeats (Figure 4). 100ng of each plasmid was taken for MDA reaction, and after 3 hours of reaction, g-TUBE was used to construct the library. BI analysis showed that the “X7” plasmid was a forward repeat sequence with a library concentration of 19ng/μL after MDA reaction. The total number of reads was 515, of which 316 were matched, accounting for 61.36%. The “X8” plasmid was a reverse repeat sequence with a library concentration of 16ng/μL after MDA reaction. The total number of reads was 769, of which 428 were matched, accounting for 55.66% (Table 7). When compared with the original sequence, deletion, insertion, and mutation occurred in both plasmids (Figure 5). It can be concluded that phi29 polymerase has a high amplification ability and strong fidelity to repeat fragments, and does not destroy the repeat structure, thus ensuring the accuracy of PacBio sequencing.

**Figure 4.**
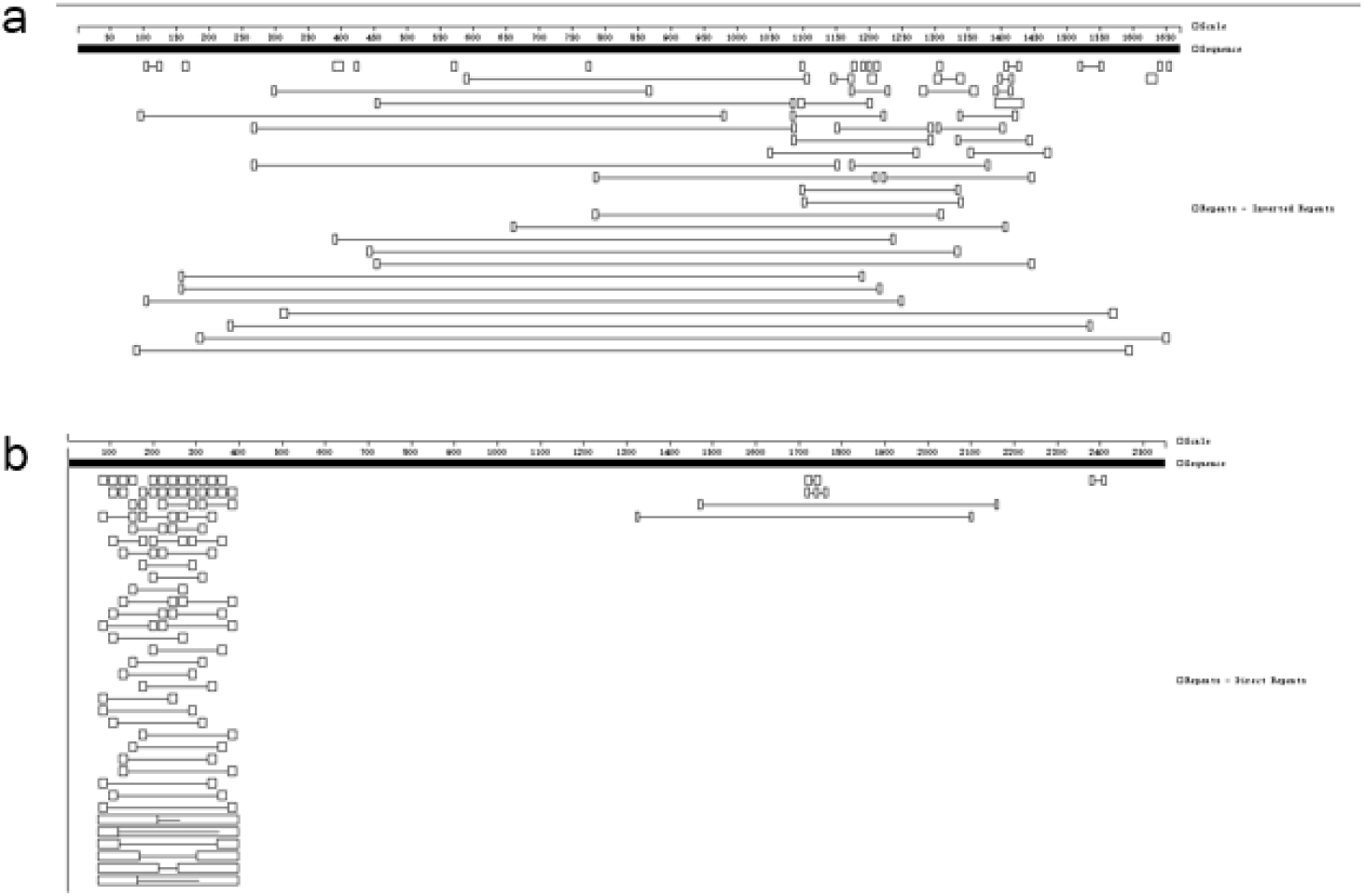
Sequence structure analysis of plasmids. The sequence structure was analyzed by Gene Quest software. (A) Direct repeat sequence structure analysis. (B)Inverted repeat sequence structure analysis.

**Figure 5.**
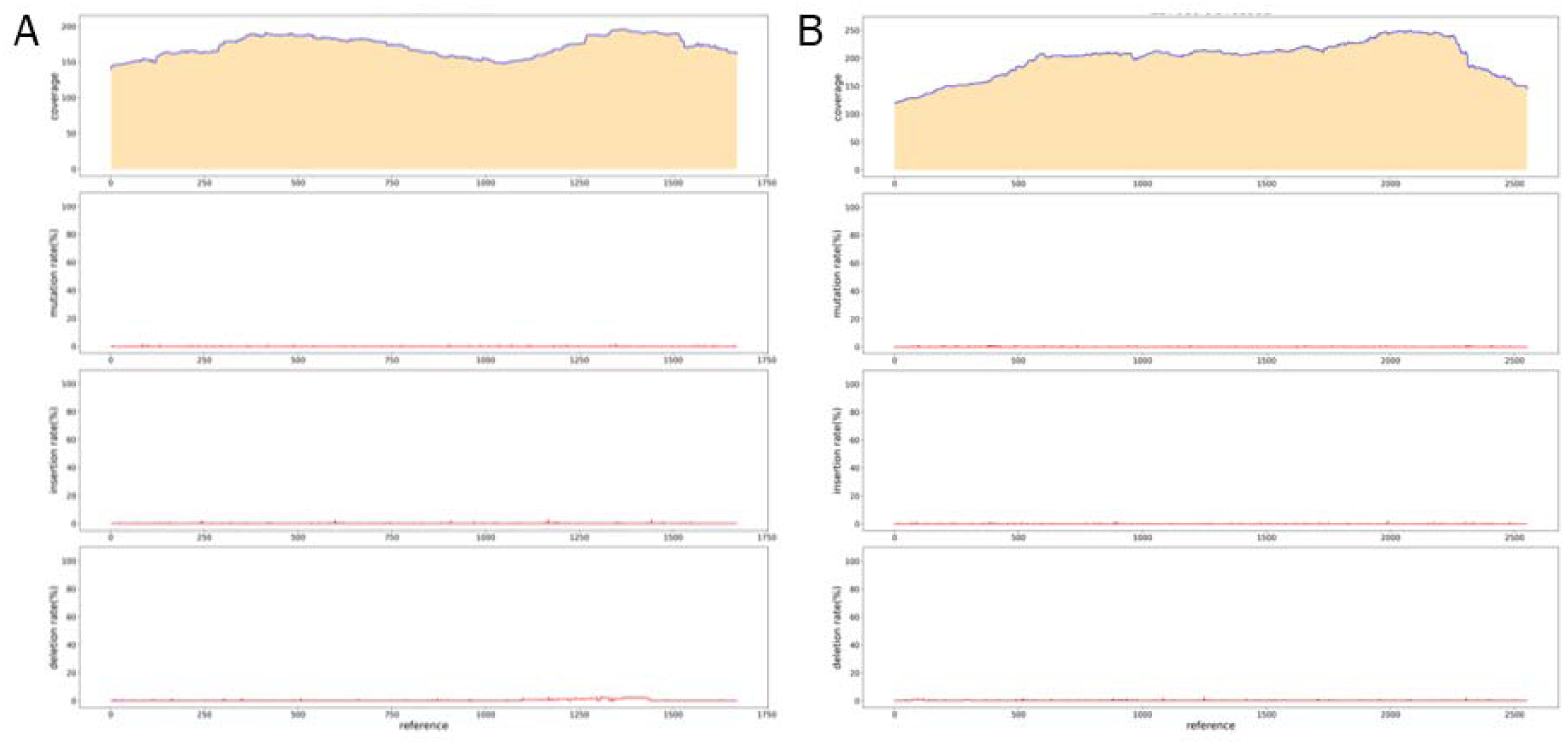
After rolling out replication, PacBio sequencing results. A. Results of PacBio sequencing of X7 plasmid. B. Results of PacBio sequencing of X8 plasmid.

**Table 7.** The amount of PacBio data used for database construction after MDA reaction.

| Sample Name | Library Concentration ( 10μL ) | Total Reads | Mapped Reads | Percent (%) | Unmapped Reads | Percent (%) |
| --- | --- | --- | --- | --- | --- | --- |
| X7 | 19 | 515 | 316 | 61.36 | 199 | 38.64 |
| X8 | 16 | 769 | 428 | 55.66 | 341 | 44.34 |

### 5. Sequencing results of PacBio after MDA of Liquid culture of bacteria contained low copy plasmid

PacBio sequencing can be performed after MDA reaction of low-copy plasmids. Therefore, we further speculate whether bacterial solution containing low-copy plasmids can be directly subjected to PacBio sequencing. Using bacterial solution sequencing can reduce the steps of plasmid extraction and thus reduce the sequencing cost. We constructed a low-copy plasmid for E. coli transformation, selected two monoclonals, and cultured them in LB medium at 37°C for 4 hours for MDA reaction. After reaction of 1μL bacterial solution for 3 hours, purified by beads, and a library was built after g-TUBE was interrupted. The concentration of DNA libraries can be reach to 24ng/ul, as shown in Table 8. After PacBio sequencing, we found that the total reads of “X9” were 434ccs, but after analysis, 66.13% were E. coli genomes. The total reads of “X10” were 414ccs, 41.3% were E. coli genomes (Table 9). It was found that a part of Escherichia coli genome was involved in MDA reaction, as long as a certain amount of data was obtained, this Escherichia coli genome would not affect PacBio sequencing. By reads number analysis, the sequencing results of the two samples were consistent with the reference sequence (Figure 6).

**Figure 6.**
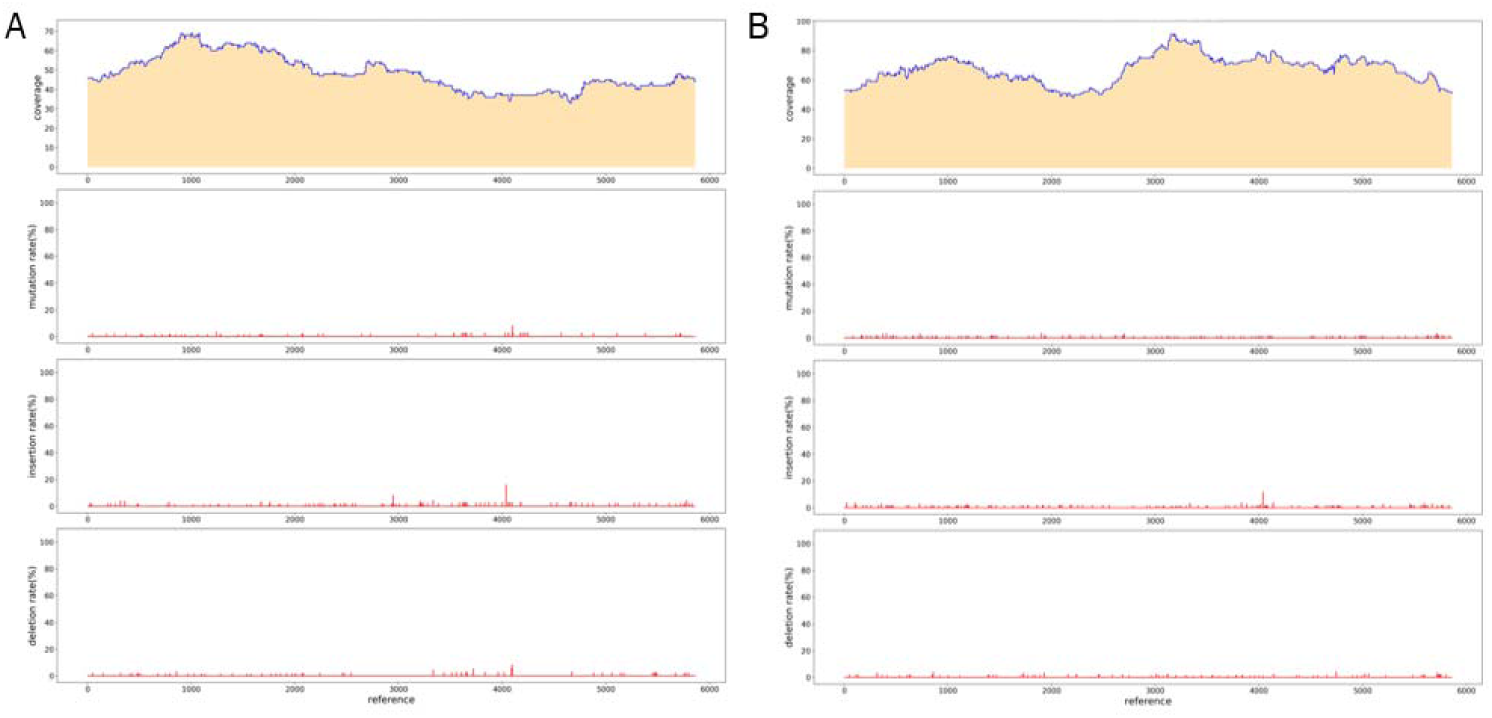
After rolling out replication, PacBio sequencing results. A. Results of PacBio sequencing of X9. B. Results of PacBio sequencing of X10.

**Table 8.** The concentration of the reservoir after MDA reaction.

| Sample Name | Library Concentration ( 10μL ) |
| --- | --- |
| X9-TP-3-1μL | 15.7 |
| X10-TP-12-1μL | 24 |

**Table 9.** The amount of PacBio data used for database construction after MDA reaction.

| Sample Name | Total Reads | Mapped Reads | Percent (%) | Unmapped Reads | Percent (%) |
| --- | --- | --- | --- | --- | --- |
| X9 | 434 | 147 | 33.87 | 287 | 66.13 |
| X10 | 414 | 243 | 58.7 | 171 | 41.3 |

## Discussion

Molecular biologists typically use circular plasmids that multiply in E. coli cells for gene cloning, expression, sequencing, and mutagenesis. These plasmids are typically divided into three groups - high-copy plasmids (equivalent to hundreds of plasmid DNA molecules per cell), low-copy plasmids (typically fewer than 50 plasmids per cell), and single-copy plasmids (about one per cell). Most of the vectors used for cloning simple DNA fragments are high copy number plasmids, which can be extracted from Escherichia coli cells with high yield. In contrast, many of the vectors used to express proteins are low-copy plasmids, and reducing the copy number (lowering the gene dose) can reduce intracellular levels of basal or non-inducible expressed proteins, which is advantageous because some foreign proteins are toxic even at very low levels and can inhibit the growth of E. coli cells, leading to difficulties in preparation.

The key to PacBio sequencing of low-copy plasmids is how to increase the yield of the library. We use the MDA reaction to increase the yield of the library by increasing the production of DNA. In general, PacBio sequencing requires the extraction of a certain amount of plasmids from E. coli. It is difficult to extract low-copy plasmids. If the plasmid transferred into Escherichia coli is mutated or missing, the low-copy plasmids in the wrong bacterial solution are extracted during sequencing, which will greatly increase the time and cost of construction. If the plasmid in the bacterial solution is verified first, without the need to extract the plasmid first, this can reduce the time and money cost of verification to a certain extent. In this experiment, we not only solved the problem of sequencing the low-copy, structured plasmid PacBio, but also established the method of sequencing PacBio without pumping the plasmid after MDA reaction with bacterial solution directly, which is an innovative technological progress.

Multiple Displacement amplification (MDA) using phage DNA polymerase phi29 isothermal amplification is a highly efficient WGA technique developed in recent years. The main challenge of the WGA approach is to obtain balanced and faithful replication of all chromosomal regions without losing or prioritizing amplification of any genomic loci or alleles. In multiple comparisons with other WGA methods, MDA appears to be the most reliable genotyping method, with the most favorable call rate, the best genome coverage, and the lowest amplification bias. In the process of sample processing, base mismatch is likely to occur, resulting in non-existent SNPS. In order to reduce the occurrence of sequencing errors, it is necessary to select a polymerase with strong fidelity, and phi29 has a strong 3 ’-5’ exonuclease reading function, and the synthesized DNA fragments have high fidelity. The exonuclease activity of this enzyme is strong, so the primers need 3 ’end thiomodification during synthesis to reduce the cutting effect of exonuclease activity on the primers. At the same time, it has good amplification ability to repeat sequences, which ensures the universality of MDA in PacBio sequencing. However, MDA still has some defects. MDA reaction will lead to the loss of poly structure and sequencing error.

## Methods

### Plasmid construction

Primers were designed to synthesize the target fragment, and the target sequence (GENEWIZ GsmartI Pfu DNA Polymerase DP1701S) was amplified by PCR. Then the vector was linearized by restriction endonuclease. Then T4 ligase (GENEWIZ T4 DNA Ligase CR1201S) was used to connect the target fragment to the carrier. Monoclone was obtained by DH5α Chemically Competent CellDL1001S (Weidibio DH5α Chemically Competent Cell DL1001). The single clone was selected and verified by pcr (GENEWIZ Gsmart Taq DNA Polymerase DP1702S), and the correct positive clone was obtained. Plasmid extraction (AXYGEN AxyPrep plasmid DNA small dose kit AP-MN-P-50) for PacBio sequencing. Multiple displacement amplification(MDA)

Before the construction of the library, mad reaction was performed to take 1μL DNA or appropriate bacterial solution for MDA amplification reaction, and 100 μM random primer was added (Invitrogen Random Primers 48190-011); 10XPhi29 MAX DNA Polymerase Reaction Buffer was supplemented with Nuclease-free ddH2O to 8.5μL, incubated at 95°C for 3 min, and immediately placed on ice to cool for 5 min. Then 10 mM dNTP was added and 1μL Phi29 MAX DNA Polymerase ( Vazyme Phi29 MAX DNA Polymerase N106-01) was incubated at 30°C for an appropriate time, and incubated at 65°C for 10 min. Magnetic bead recovery(Vazyme VAHTS DNA Clean Beads N411-01)。

### PacBio library database preparation

The plasmid was interrupted with g-TUBE (Covaris g-TUBE 520079), and the plasmid was repaired with New England Biolabs PreCR® Repair MixM0309L. 1μL of 5: 1dATP (Vazyme dATPP034-01)/dNTP (Vazyme dNTP Mix P031-01); 0.5μL T4 PNK (Vazyme T4 Polynucleotide KinaseN102-01); 2μL T4 DNA Polymerase (Vazyme T4 DNA polymerase N101-01); 0.5μL rTaq (Takara TaKaRa Taq™ R001A) plus A tail for connection, 8.0 μLPrD (aladdin PrD P103430) at the end; 10.0 μL50% PEG-8000 (solarbio PEG 8000 25322-68-3); 4.0μLT4 DNA Ligase (Vazyme T4 DNA Ligase C301-01) was connected to the interrupted plasmid, the library was constructed, and the fragment length was tested using Agilent 2100 for quality inspection. The constructed SMRTbell library, combined with the upper primers and polymerase of the SMRTbell Express Template Prep Kit 2.0 kit, was added to the sequencing chip of the PacBio platform in a freely diffused manner for sequencing.

### PacBio sequencing data analysis

The original disembarking data is first corrected by using ccs (4.2.0) [8] software for subread generated by the same zero-mode waveguide hole (ZMW) to obtain a high-quality CCS sequence. minimap2 (version 2.15-r905) [9] software is used to compare the high-quality CCS sequence to the reference sequence. Finally, samtools(Version: 1.9) [10] software was used for quality control of the comparison results. Specific quality control parameters include the minimum cycle sequencing times, the minimum prediction accuracy, the minimum insertion fragment length and the maximum insertion fragment length, and the set values of the above parameters are 5,90,1 400 and 1 800, respectively. After quality control, the qualified sequences were divided into different samples according to barcode sequence. Meanwhile, barcode and primer information of each sequence were deleted for subsequent analysis.

## ACKNOWLEDGMENTS

We thank members of Azenta life science for their technical support.

## Supplement

S1, S3, S5, S7 Tabl : Results of PacBio sequencing of X1, X2, X3 and X4 plasmid. The plasmid was not treated with MDA reaction.

S2, S4 Tabl : Results of PacBio sequencing of X1-MDA and X2-MDA plasmid. MDA reaction was performed before building the library.

S6, S8 Tabl : Results of PacBio sequencing of X3-MDA and X4-MDA plasmid. MDA reaction was performed after building the library.

S9, S10 Tabl : Low-copy plasmids PacBio sequencing results

S11 Tabl : Forward repeated structure PacBio sequencing results S12 Tabl : Intensive repeated structure PacBio sequencing results

S13, S14 Tabl : Results of PacBio sequencing of X9 and X10.The MDA reaction was performed on E. coli.

